# No objective evidence of neuropsychological deficits in people with subjective cognitive changes following COVID-19 infection

**DOI:** 10.64898/2026.06.03.723612

**Authors:** Samantha Abram, Kaitlyn Dal Bon, Daniel H. Mathalon, Susanna Fryer, Sara Song, Zanib Naeem, Norina Tang, Lynn Pulliam, Judith M. Ford

## Abstract

**Background:** Among those who develop Long COVID, many experience persistent cognitive “brain fog.” The degree to which these subjective complaints reflect measurable neuropsychological deficits remains unclear. Prior work suggests that subjective cognitive impairment may be more closely associated with affective symptoms than objective performance. This study examines the relationship between subjective and objective cognitive function in adults with post-COVID cognitive complaints, and assesses the association between self-reported deficits and biological markers of dementia risk and inflammation.

**Methods:** Eighty-six adults with prior COVID-19 infection (mean age 41.5 years) completed neuropsychological testing (MATRICS; CVLT-III) and self-report measures of depression and anxiety (Beck Inventories). Participants were classified as *Cases* (*n*=47) if they endorsed worsening memory or concentration since infection or *Controls* (*n*=38) if they did not. Objective cognitive impairment was defined as performance <1 SD below normative means on at least one test. APOE-ε4 status and soluble CD14 (sCD14) levels were assessed.

**Results:** Cases reported higher depression and anxiety symptoms than Controls (both *p*<0.001), but groups did not differ on objective cognitive performance (*p*=0.39). Cases were more likely to be APOE-ε4 carriers (*p*=0.01) and had higher sCD14 levels (*p*=0.01). Neither marker was associated with objective performance.

**Conclusions:** Subjective cognitive complaints following COVID-19 were not accompanied by measurable neuropsychological deficits but were linked to elevated affective symptoms and biological risk markers. Findings highlight a dissociation between perceived and objective cognition and suggest that inflammatory, genetic, and affective factors may shape self-perceived decline. Longitudinal studies are needed to determine whether these markers confer vulnerability to cognitive decline.

## 1. INTRODUCTION

There is established evidence that COVID-19 is associated with a wide spectrum of acute neurological and psychiatric complications (Beghi et al., 2022; Ellul et al., 2020; Varatharaj et al., 2020). Taquet et al. (2021) estimated that the incidence of a neurological or psychiatric diagnosis within six months after infection was 33.6%. Individuals with post-COVID neurological or psychiatric complications were also more likely to experience impairments in daily living and employment, as well as depressive symptoms (Shil et al., 2025). Elboraay et al. (2025) similarly reported high rates of persistent cognitive and affective complaints, including memory disorders (27.8%), cognitive impairment (27.1%), concentration impairment (27.1%), depression (14.0%), and anxiety (13.2%). Given these widespread and persistent symptoms, clarifying how individuals *perceive* their cognitive changes versus how they *perform* on objective measures is critical for understanding the mechanisms and clinical implications of post-COVID cognitive dysfunction.

Tarantini et al. (2025) first addressed whether mood and subjective experience of cognitive decline correspond with objective cognitive deficits following COVID-19. Using standardized tests of attention, memory, and executive function, they found that subjective cognitive complaints were more strongly associated with depressive symptoms than with objective cognitive performance, underscoring that self-perceived deficits—how well you *think* you are doing—may not reliably reflect actual cognitive ability (Burmester et al., 2016). Objective cognitive impairment was identified in only 21% of patients on tests of attention and 18% on tests of memory and executive function, with 39% showing impairment in at least one domain. These findings highlight the need to evaluate both subjective and objective cognitive outcomes, particularly in a condition such as COVID-19 where affective symptoms may shape symptom perception.

Standardized neuropsychological tests remain essential for distinguishing between self-reported concerns and measurable deficits. These assessments are widely used across neurological and psychiatric conditions to detect subtle impairments, monitor progression, and evaluate treatment response. In the context of COVID-19, subjective cognitive complaints may be influenced by depression, anxiety, or fatigue; thus, pairing objective tests with validated self-report instruments of mood, such as the Beck Depression Inventory (BDI) and Beck Anxiety Inventory (BAI), is necessary for interpreting the origins and significance of perceived decline.

Beyond psychological factors, biological risk markers may help identify individuals more vulnerable to post-COVID cognitive dysfunction. The APOE-ε4 allele, a well-established genetic risk factor for Alzheimer’s disease, has been associated with increased susceptibility to neuroinflammation and cognitive decline in several post-infectious contexts (Dias et al., 2025). This risk marker also corresponds with cognitive impairment among non-demented healthy aging APOE-ε4 carriers (Small et al., 2004). In a sample of people following COVID-19 infection, APOE-ε4 carriers exhibit significantly poorer performance in executive functioning on neuropsychological tests, and this effect was independent of subjective cognitive complaint (Tarantini et al., 2025). Elevated levels of the inflammatory monocyte activation marker, soluble CD14 (sCD14), was previously linked to subjective cognitive complaints in this cohort (Pulliam et al., 2024; Tang et al., 2025). sCD14 is elevated in people with several acute viral infections including COVID-19 (Martin et al., 2020; Zingaropoli et al., 2021) and sCD14 levels predict COVID-19 disease severity (Sharygin et al., 2023). Introducing APOE-ε4 and sCD14 markers allows for evaluations of whether subjective symptoms, objective cognitive performance, or mood disturbances may be partially driven by immunological or genetic vulnerability.

Here we examine the similarities between subjective and objective cognitive decline by using standardized neuropsychological assessments and self-reported changes in function and mood from pre-infection state. Our study focused on individuals who solely received outpatient treatment for COVID-19 infection and were not hospitalized. We hypothesized that individuals reporting persistent sequelae would show poorer performance on standardized neuropsychological tests, with cognitive scores at least one standard deviation below the normative mean, consistent with the metric used by Tarantini et al. (2025). We further predicted that these individuals would report elevated symptoms of depression and anxiety, as measured by the BDI and BAI. In addition, we hypothesized that higher BDI and BAI scores would be associated with increased levels of inflammatory markers (i.e., sCD14), given previous findings in this cohort. Finally, we tested whether objective measures of brain function were related to both APOE-ε4 status and inflammatory markers to evaluate whether genetic and immunological factors jointly contribute to cognitive and affective outcomes.

## 2. MATERIALS AND METHODS

### Sample Demographics

We report data from a total of 86 males (47.7%) and females who had experienced COVID-19, confirmed by a positive viral RNA PCR or antigen test result from nasal or throat swab, more than 12 weeks prior (Table 1). Participants were recruited from the San Francisco Veterans Affairs Health Care System and San Francisco Bay Area and were required to be proficient in English. Exclusion criteria included a self-reported history of head injury, neurological disorders, or other significant medical condition affecting the central nervous system. Urine toxicology screening for common drugs of abuse (e.g., opiates, cocaine, amphetamines, phencyclidine) was conducted, and individuals with positive results were excluded. To reduce potential confounds related to hospital-based interventions for COVID-19 that may affect cognition (such as intubation or mechanical ventilation), participants with a history of COVID-19-related hospitalization were excluded. Self-reported information about previous and current psychiatric diagnoses was recorded. Written informed consent was obtained from all participants following a detailed explanation of the study procedures. The study was approved by the Institutional Review Boards at the University of California, San Francisco, and the San Francisco Veterans Affairs Medical Center (SFVAMC).

**Table 1.**
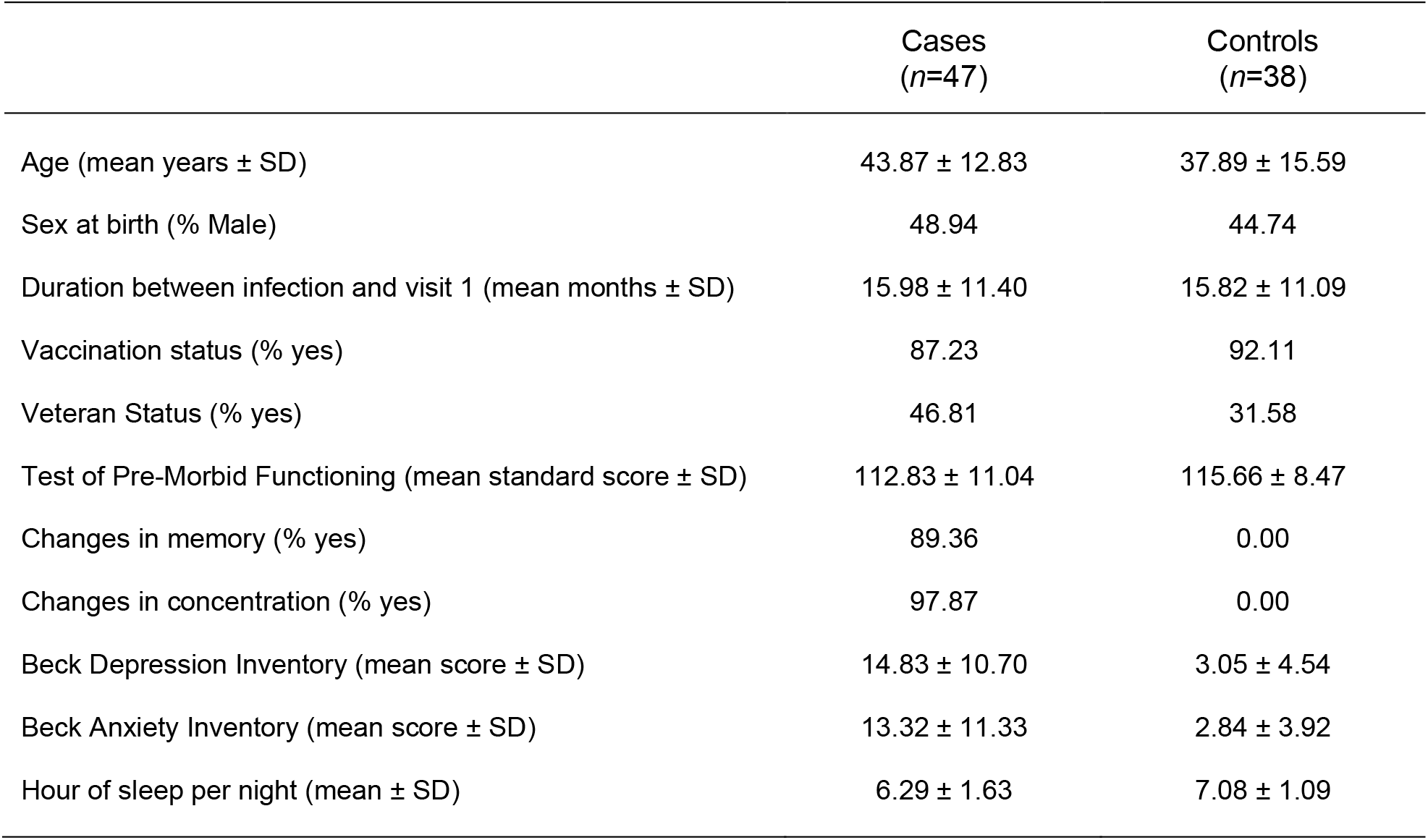
Demographic Data & Clinical Characteristics of Participant Groups.

### Subjective Cognitive Change (Self Report)

Cases were defined as individuals who self-reported worsening of memory or concentration (*n* = 47) following infection. Controls were defined as those who did not endorse worsening in either domain (*n* = 38). Thus, subjective cognitive change was a dichotomous variable where 1 = Cases, 0 = Controls. Memory and concentration complaints were highly intercorrelated (r =.792, *p* < 0.001). Cases were more likely to have a prior diagnosis of anxiety (Pearson Chi-Square = 5.23, *p* = 0.022) and/or depression (Chi-Square = 4.60, *p* = 0.032), with a similar trend for PTSD (Chi-Square = 3.71, *p* = 0.054). Cases were slightly older than Controls at a trend level but did not differ by vaccination status, veteran status, pre-morbid functioning, or biological sex (Table 1).

### Objective Cognitive Impairment (Neuropsychological Assessment)

Scores from the Measurement and Treatment Research to Improve Cognition in Schizophrenia (MATRICS) Consensus Cognitive Battery (Nuechterlein et al., 2008) and California Verbal Learning Test-Version III (CVLT-III; Delis et al., 2017) were used to derive domain scores for the following: speed of processing (BACS: Symbol Coding, Category Fluency: Animal Naming, Trail Making Test: Part A), attention/vigilance (Continuous Performance Test – Identical Pairs), working memory (Letter-Number Span), reasoning and problem-solving (NAB: Mazes), and verbal learning (Immediate and Delayed Recall). Scores were standardized across the normal population. We defined objective cognitive impairment as scoring one standard deviation below the mean (T-score < 40 or Scaled Score < 7) on speed of processing, attention/vigilance, working memory, reasoning and problem-solving, or verbal learning; this yielded a dichotomous variable where 1 = presence of objective cognitive impairment, 0 = absence of objective cognitive impairment. This approach is comparable to the aforementioned report by Tarantini et al. (2025). Our primary analyses used this composite cognitive impairment metric; for completeness, we also report the distributions and relevant statistics for individual subtests in Table 2.

**Table 2.**
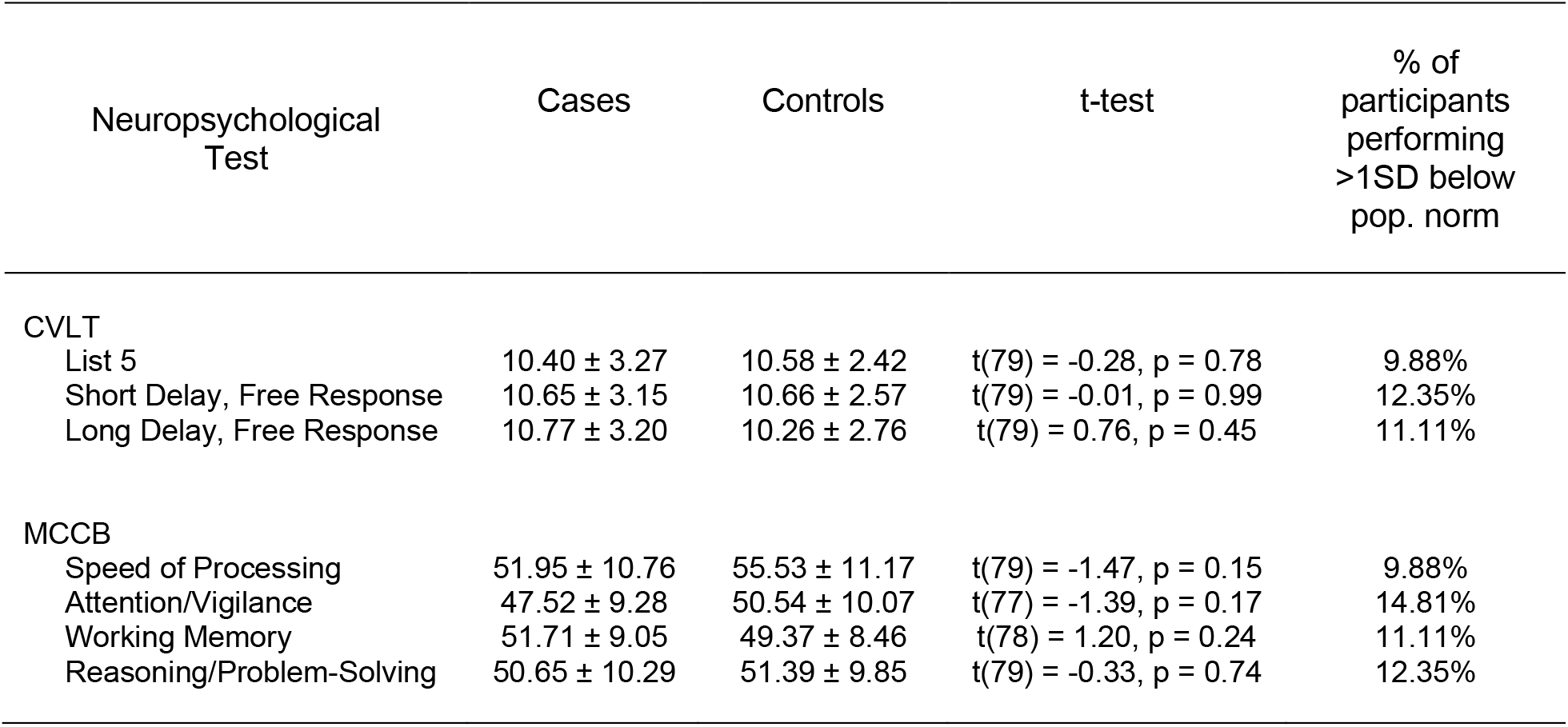
Neuropsychological Test Outcomes (mean ± SD)

### Assessment of Current Depression and Anxiety

We used the self-report Beck Depression Inventory (BDI) and the Beck Anxiety Inventory (BAI) to assess current symptoms of depression and anxiety (Beck et al., 1988, 1996).

### APOE-ε4 Genotyping and sCD14 Quantification

These were performed as previously published on these cohorts (Tang et al., 2024, 2025).

### Statistical analysis

To test relations between subjective cognitive change and psychiatric symptoms, we fit two linear regression models predicting current depressive and anxiety symptom severity from self-reported cognitive change, including age and a cognitive change × age interaction terms (i.e., Depression ∼ Subjective Cognitive Change × Age; Anxiety ∼ Subjective Cognitive Change × Age).

We next used a binomial regression model to test the association between objective cognitive impairment and subjective cognitive change, controlling for age (i.e., Objective Cognitive Impairment ∼ Subjective Cognitive Change × Age).

To examine biological and inflammatory risk markers, we fit four additional binomial or linear regression models regressing APOE-ε4 status (binary) or sCD14 levels (continuous) on objective cognitive impairment or subjective cognitive change following COVID-19 infection, with age interaction terms included (i.e., APOE-ε4 ∼ Cognitive Impairment × Age; APOE-ε4 ∼ Subjective Cognitive Change × Age; sCD14 ∼ Cognitive Impairment × Age; sCD14 ∼ Subjective Cognitive Change × Age).

For all models, interaction terms were tested and removed when nonsignificant, yielding additive models with a common age slope. False discovery rate (FDR) correction was applied to adjust for multiple comparisons within related model families (e.g., two psychological variable models, four biological/inflammatory risk marker models).

## 3. RESULTS

### Cases report greater current depression and anxiety symptoms

Cases reported greater depression (*t*_82_ = 6.33, *p* < 0.001, *p*_adj_ < 0.001) and anxiety (*t*_82_ = 5.42, *p* < 0.001, *p*_adj_ < 0.001) than Controls (Figure 1). The age interaction was not significant, and there was no main effect of age after reducing to a common slope model.

**Figure 1.**
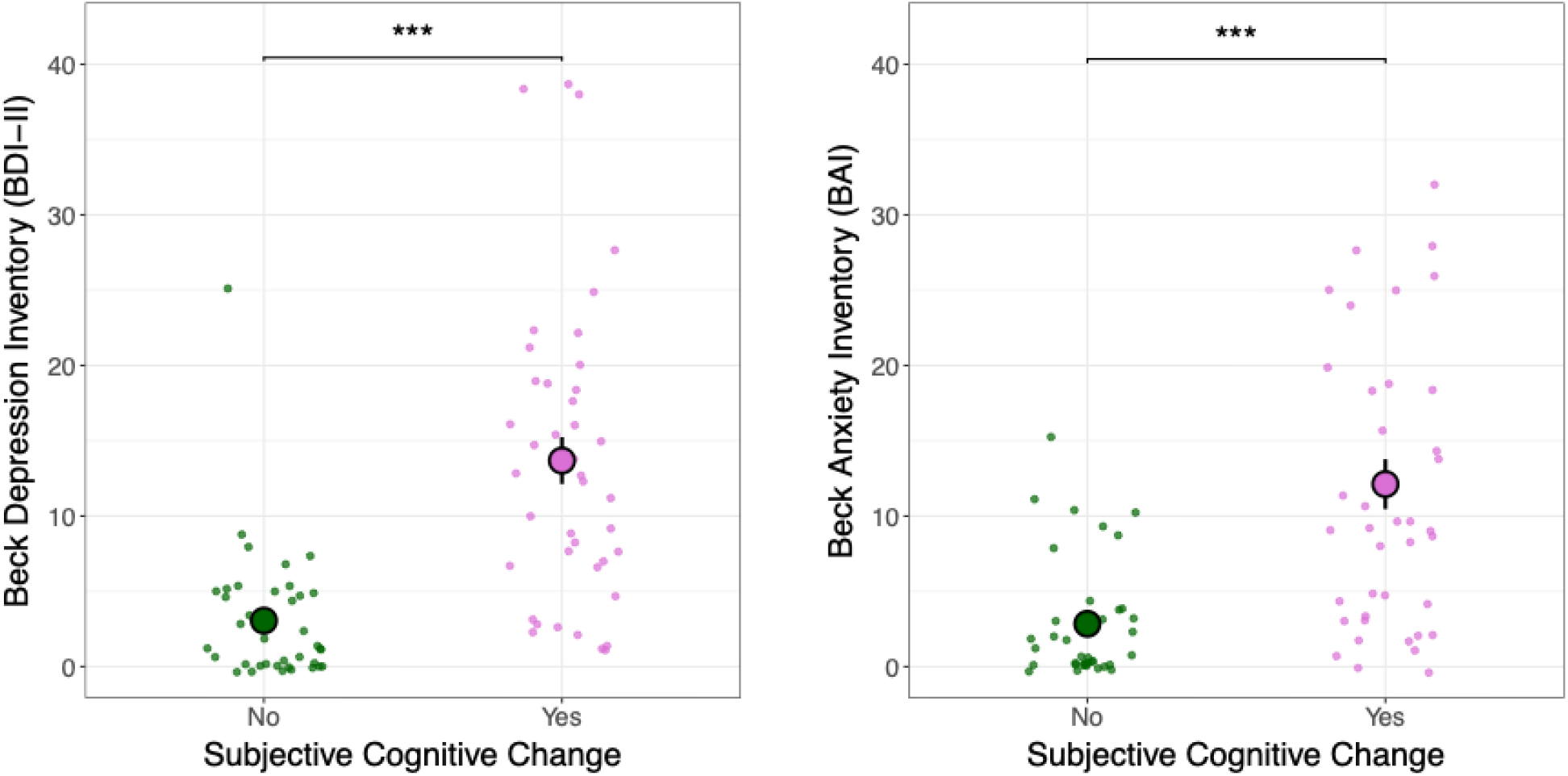
People who reported subjective cognitive change after COVID-19 infection also report higher depression (*left*) and anxiety (*right*) symptoms. \*\*\**p* < 0.001.

### Cases do not perform worse on objective measures of cognition than Controls

Using the binomial regression model described in the Methods, Cases and Controls did not differ from each other in objective cognitive impairment. The main effect of Group (e.g., presence vs. absence of subjective cognitive change) was not significant (*z*_76_ = 0.86, *p* = 0.39; Figure 2), indicating no association between subjective complaints and objective cognitive performance. This pattern suggests dissociation between subjective and objective reports of cognitive functioning. The age interaction was not significant, and age was not a significant predictor after reducing to a common slope model.

**Figure 2.**
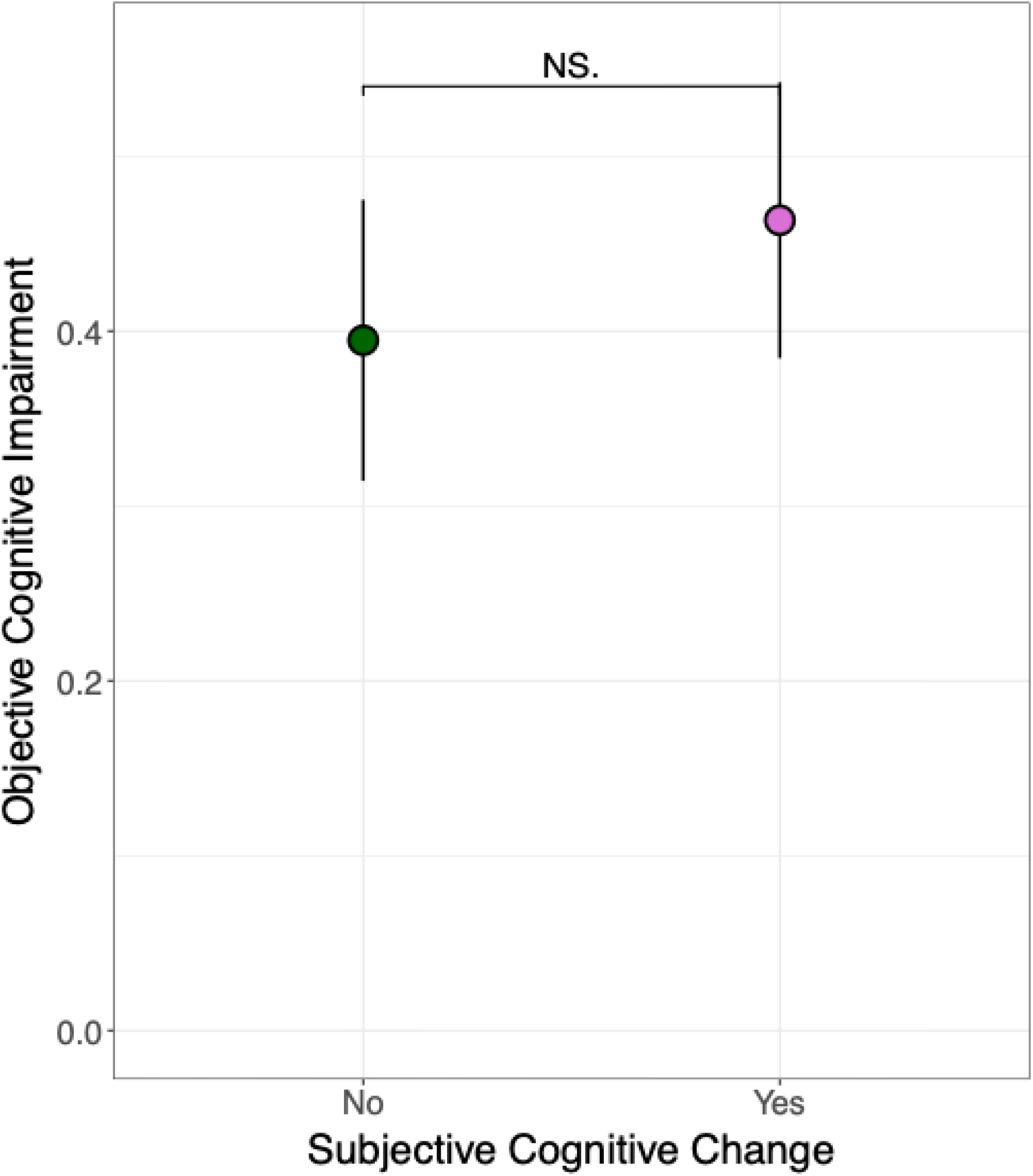
People who reported subjective cognitive change after COVID-19 did not show evidence of cognitive impairment on standardized neuropsychological tests. The y-axis indicates proportion of participants considered to have objective cognitive impairment (i.e., 34% of Controls and 44% of Cases had objective cognitive impairment).

Table 2 shows that there were no significant differences between Cases and Controls on any individual neuropsychological subtest. And for any given subtest, about 12% or fewer of participants, across the whole sample, were considered to have cognitive impairment in that domain; as defined by performance more than 1 SD below the population mean of 50 for the MCCB and 7 for the CVLT (Table 2). We also note that comparable non-significant Case versus Control effects were observed when using a continuous objective cognitive functioning variable that averaged across the seven neurocognitive domains.

### Cases have higher rates of biological markers associated with dementia risk and inflammation

Cases had higher rates of APOE-ε4 status (*t*_82_ = 2.49, *p* = 0.01, *p*_adj_ = 0.03) (Figure 3; Tang et al., 2025); the age interaction and common slope terms were both not significant. However, we did observe a Group × Age effect for sCD14 protein levels (*t*_81_ = -2.08, *p* = 0.04, *p*_adj_ = 0.08) that remained at a trend-level of significance after multiple comparison correction (Figure 3). Follow-up analyses revealed a positive age correlation in Controls only (B = .04, *t*_81_ = 3.79, *p* < .001) and a null slope in Cases (*p* = .52). Furthermore, higher sCD14 proteins among Cases were particularly evident when comparing younger participants (*t*_26_ = -3.78, *p* <. 001). If we were to drop the age interaction term based on it not surviving multiple comparison correction, we still observe a significant Cases versus Controls effect such that Cases have higher sCD14 on average (*t*_82_ = 2.64, *p* = 0.01, *p*_adj_ = 0.03).

**Figure 3.**
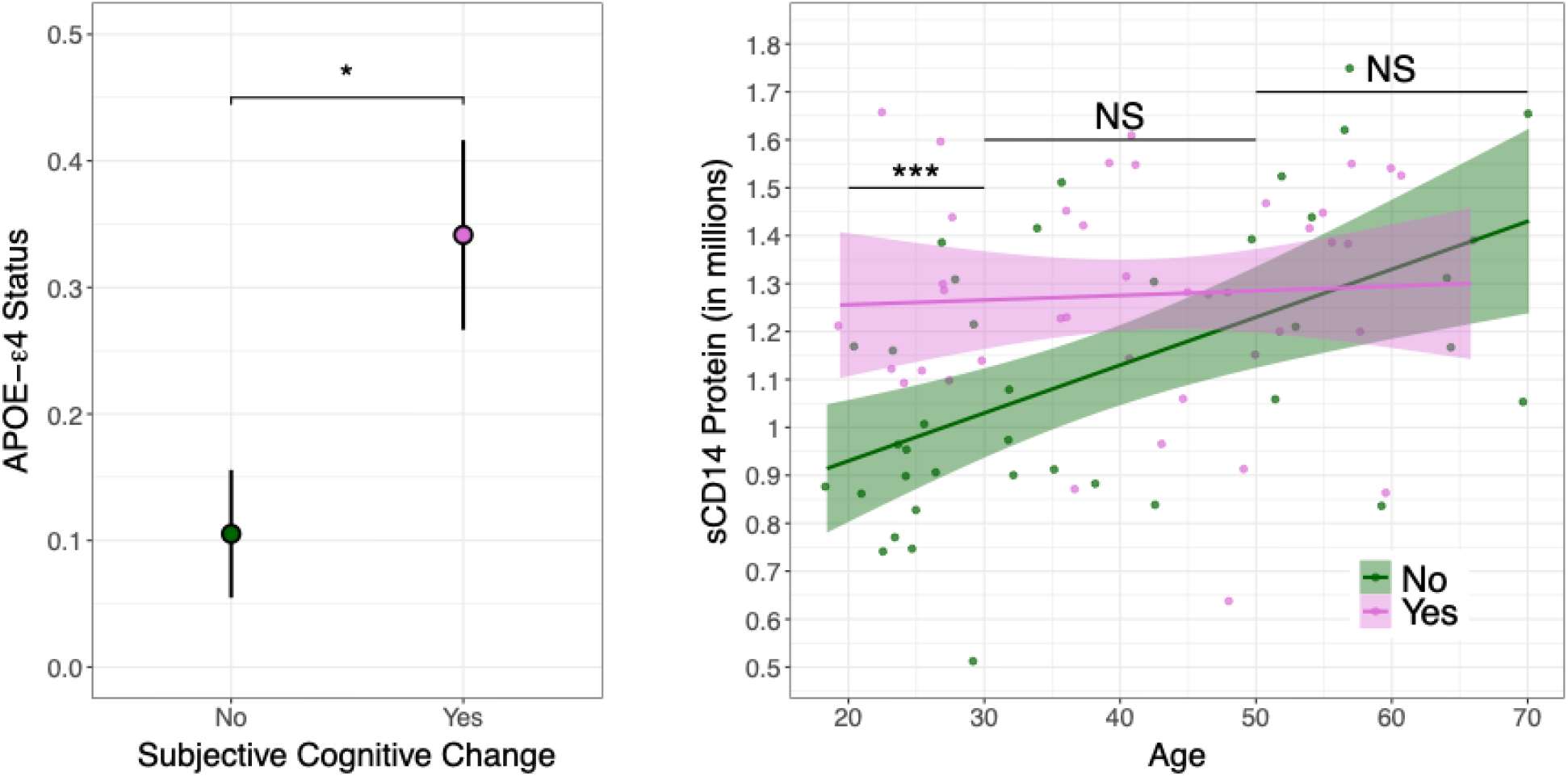
People who reported subjective cognitive change after COVID-19 were more likely to be APOE-ε4 positive (*left*). The y-axis indicates proportion of participants with a positive APOE-ε4 status (i.e., 0.11% of Controls and 0.36% of Cases had positive APOE-ε4 status). \**p* < 0.05; \*\*\**p* < 0.001. Higher sCD14 protein levels were observed in participants who endorsed subjective change, particularly when comparing younger participants (*right*).

Neither APOE-ε4 nor sCD14 correlated directly with objective measures of cognition (both *ps*_unadjusted_ > 0.6). Both APOE-ε4 and sCD14 remained significant when included in the same model predicting self-reported subjective cognitive change (both *ps* < 0.05). Current depressive and anxiety symptoms correlated directly with APOE-ε4 and sCD14 (all *ps* < 0.05).

A subset of people had data on additional blood markers that have been linked with COVID-19, including the pro-inflammatory cytokine, Interleukin-6 (IL-6), and C-reactive protein (CRP) (Tang et al., 2024; Tarantini et al., 2025). IL-6 was unrelated to subjective memory changes (*p* = 0.77) but was positively associated with age (*t*_41_ = 3.21, *p* = 0.002). CRP was not correlated with subject memory changes or age (both *ps* > 0.7). The age interaction terms were not significant for either IL-6 or CRP. Neither IL-6 nor CRP were associated with depression or anxiety level (all *ps* > 0.6).

## 4. Discussion

We found no evidence of objective neuropsychological deficits in people reporting subjective cognitive decline following COVID-19 infection. Like Tarantini et al. (2025), we observed elevated levels of self-reported depression and anxiety in Cases compared to Controls. We then expanded their findings by associating subjective cognitive complaints with markers of inflammation (sCD14) and dementia risk factor (APOE-ε4).

Participants in our study who reported worsening cognition following COVID-19 infection did not perform worse on standard neuropsychological tests. As noted previously, our *subjective* and *objective* evaluations of our own functioning do not always align. In a systematic review and meta-analysis, Burmester et al. (2016) asked about the relationship between subjective and objective cognitive complaints in aging. The authors concluded that comprehensive measures of subjective cognitive complaints were associated independently with both objective cognitive function and depressive symptoms. Our findings extend this literature by showing that only subjective cognitive complaints are associated with depression, whereas objective cognitive impairment is not associated with either outcome. We posit a few considerations for this null finding: we asked if people thought they were doing worse than before COVID-19 infection, but we did not ask them if they thought they were impaired compared to others. We do not know how the participants in our study would have scored before COVID; we only know how someone compares to population norms. Additionally, a drop in memory performance may not land someone in the abnormal range if they have adequate cognitive reserve. As a group, participants in this study were not cognitively impaired, and the proportions of participants classified as impaired on individual subtests were lower than those reported by Tarantini (e.g., about 10% versus 20% across the various subtests). It is therefore possible that a study including a more impaired sample would reveal a stronger effect of objective cognitive impairment. Another distinction is that none of our patients were hospitalized for COVID-19 infection, and could therefore represent a lower severity group.

Cases scored higher on self-report tests of depression and anxiety than Controls in our study. There are high rates of depression and anxiety reported in people who endorse long COVID-19 symptoms (Fancourt et al., 2023; Graham et al., 2021; Renaud-Charest et al., 2021). Depression is associated with negative self-assessments (Schweizer et al., 2018) or less positive self-assessments, than in non-depressed people (Pham, 2007). One explanation is that individuals self-reporting cognitive impairments are more self-critical.

Depression symptoms may also represent either a risk factor for, or an early prodromal indicator of, dementia (Brommelhoff et al., 2009; Wiels et al., 2020). Supporting the prodromal hypothesis, depression and anxiety symptoms have been linked to APOE-ε4 status, a genetic variant that increases risk for Alzheimer’s disease (Wang et al., 2019). At the same time, the relationship appears to be bidirectional: subjective cognitive decline can itself be distressing and may contribute to or exacerbate depressive symptoms. In this sense, reduced cognitive reserve and emerging cognitive difficulties may heighten feelings of depression, creating a bidirectional relationship between mood and cognitive complaints.

APOE-ε4 is the strongest known genetic risk factor for late-onset Alzheimer’s disease and is linked to both objective cognitive decline (Tarantini et al., 2025) and subjective cognitive complaint (Munro et al., 2023). In our sample, APOE-ε4 was associated with subjective cognitive change, but not objective cognitive impairment. Biological factors related to neurodegeneration and inflammation may contribute to perceived cognitive difficulties. Carriers may be more attuned to subtle cognitive fluctuations or experience early neurobiological changes that do not yet manifest as detectable deficits. The present findings are consistent with that pattern, suggesting that APOE-ε4 status may influence cognitive self-appraisal or interact with affective symptoms to heighten perceived decline. And similar to our point above, the lack of association with objective cognitive impairment may be due to the fact that only about 10% of this sample showed objective impairment on our neuropsychological tests.

sCD14, a marker of monocyte activation and systemic inflammation, has also been associated with neuroinflammation and risk for cognitive aging and dementia (Pase et al., 2020). Elevated sCD14 may contribute to subjective experiences such as mental fatigue or “brain fog,” which are frequently reported after COVID-19, even in the absence of measurable decrements on standardized neuropsychological tests. In the current sample, we observed a trend level age by group interaction, such that among Controls, sCD14 demonstrated a positive association with age, whereas Cases exhibited elevated sCD14 levels regardless of age. This pattern resulted in the largest group differences emerging among younger participants (aged 30 years or younger). One possible explanation is that younger individuals would be expected to have relatively low baseline inflammation; therefore, elevated inflammatory markers in younger Cases may reflect a more salient deviation from normative physiological levels, increasing their perceptibility and subjective impact (Furman et al., 2019). Consistent with this interpretation, sCD14 was associated with subjective cognitive complaints but not objective cognitive impairment. Together, our results suggest that inflammation- and risk-related biological factors may disproportionately influence individuals’ perception of cognitive inefficiency rather than their actual cognitive performance (Franceschi et al., 2018).

We also observed that both sCD14 levels and APOE-ε4 carrier status were associated with elevated depression and anxiety symptoms. Prior work has demonstrated robust associations between APOE-ε4, depression, and subjective cognitive decline (see meta-analysis by Huang et al., 2020), suggesting that genetic vulnerability may interact with affective and inflammatory processes to shape cognitive symptom reporting. sCD14 has similarly been linked to depressive symptoms, particularly in populations with HIV, highlighting its role as a marker of monocyte activation in post- or active viral infection contexts (Hussain et al., 2023; Veihman et al., 2025). Notably, our findings were specific to sCD14: inflammatory markers that are commonly studied in major depressive disorder, including IL-6 and CRP (Poletti et al., 2024) were neither elevated in cases nor correlated with depression or anxiety symptoms in our sample. This pattern suggests that viral-related immune activation, rather than the low-grade systemic inflammation typically reported in depression, may be more relevant to affective symptoms in this cohort. In terms of the directionality of these relationships, heightened inflammation and elevated genetic risk for dementia may contribute directly to mood symptoms and perceived cognitive impairment (Slavich & Irwin, 2014); alternatively, depression and anxiety may exacerbate subjective cognitive complaints and be accompanied by secondary inflammatory changes (Berk et al., 2013). Disentangling these pathways will require longitudinal data to clarify temporal ordering and causal mechanisms.

This study is not without limitations. As noted above, our study lacked pre-COVID baseline neurocognitive, psychiatric, or biological data. This means we cannot be sure that people reporting subjective cognitive changes did not experience a post-infection decline. We also do not know whether inflammation (i.e., sCD14 levels) was already elevated (due to psychiatric symptoms) or instead reflect a more specific post-COVID consequence. Although we asked participants if they ever had a diagnosis of anxiety or depression, we did not confirm this with a diagnostic interview to assess past depression or anxiety nor did we do a structured assessment (e.g., Structured Clinical Interview for DSM-V) at study enrollment. Further, while the Cases endorsed more depressive and anxiety symptoms on the Beck Inventories, these self-report measures do not replace clinician-guided structured interviews.

To conclude, subjective cognitive complaints following COVID-19 were not accompanied by measurable neuropsychological deficits but were linked to elevated affective symptoms and biological risk markers (APOE-ε4, sCD14). These findings highlight a dissociation between perceived and objective cognitive function in post-COVID populations and suggest that inflammatory and genetic factors may shape self-perceived cognitive decline even in the absence of impairment. Longitudinal studies are needed to determine whether these biological markers confer increased vulnerability to future cognitive decline.

## Acknowledgements

The authors have no relevant financial interests to report. SVA, KDB, DHM, SLF, ZN, NT, LP, and JMF were U.S. Government employees at the time of data collection. The content of this manuscript is solely the responsibility of the authors and does not necessarily represent the views of the Department of Veterans Affairs.

## Ethical Standards

The study was approved by the institutional review board of UCSF, and all participants provided written informed consent.

